# Genomic factors limiting the diversity of Saccharomycotina plant pathogens

**DOI:** 10.1101/2025.02.27.640420

**Authors:** Sun Lee, Caroline West, Dana A. Opulente, Marie-Claire Harrison, John F. Wolters, Xing-Xing Shen, Xiaofan Zhou, Marizeth Groenewald, Chris Todd Hittinger, Antonis Rokas, Abigail Leavitt LaBella

**Affiliations:** North Carolina Research Center (NCRC), Department of Bioinformatics and Genomics, The University of North Carolina at Charlotte, Kannapolis, NC 28081, U.S.A.; Biology Department, Villanova University, Villanova, PA 19085, U.S.A.; Department of Biological Sciences, Vanderbilt University, Nashville, TN 37235, U.S.A.; Evolutionary Studies Initiative, Vanderbilt University, Nashville, TN 37235, U.S.A.; Laboratory of Genetics, Wisconsin Energy Institute, Center for Genomic Science Innovation, J. F. Crow Institute for the Study of Evolution, University of Wisconsin–Madison, Madison, WI 53726, U.S.A.; DOE Great Lakes Bioenergy Research Center, University of Wisconsin–Madison, Madison, WI 53726, U.S.A.; Centre for Evolutionary and Organismal Biology, Institute of Insect Sciences, Zhejiang University, Hangzhou, China; Guangdong Province Key Laboratory of Microbial Signals and Disease Control, Integrative Microbiology Research Center, South China Agricultural University, Guangzhou, China; Westerdijk Fungal Biodiversity Institute, Utrecht, The Netherlands; Center for Computational Intelligence to Predict Health and Environmental Risks (CIPHER), University of North Carolina at Charlotte, Charlotte, North Carolina, U.S.A.

**Author notes:** Corresponding Author: A.L. LaBella.

**Keywords:** Saccharomycotina, fungi, phytopathogen, reverse ecology

## Abstract

The Saccharomycotina fungi have evolved to inhabit a vast diversity of habitats over their 400-million-year evolution. There are, however, only a few known fungal pathogens of plants in this subphylum, primarily belonging to the genera *Eremothecium* and *Geotrichum*. We compared the genomes of 12 plant-pathogenic Saccharomycotina strains to 360 plant-associated strains to identify features unique to the phytopathogens. Characterization of the oxylipin synthesis genes, a compound believed to be involved in *Eremothecium* pathogenicity, did not reveal any differences in gene presence within or between the plant-pathogenic and plant-associated strains. A reverse-ecological approach, however, revealed that plant pathogens lack several metabolic enzymes known to assist other phytopathogens in overcoming plant defenses. This includes L-rhamnose metabolism, formamidase and nitrilase genes. This result suggests that the Saccharomycotina plant pathogens are limited to infecting ripening fruits as they are without the necessary enzymes to degrade common phytohormones and secondary metabolites produced by plants.

## Introduction

Across the fungal kingdom, plant pathogens are highly diverse, with a concentration in the phyla Ascomycota and Basidiomycota (Doehlemann et al. 2017). Despite containing over 1,000 species that have evolved over the past 400 million years (Opulente et al. 2024) the subphylum Saccharomycotina, which belongs to the Ascomycota, contains only one well-studied plant pathogenic genus: *Eremothecium.* Recent work has shown, however, that of 1,088 Saccharomycotina strains examined, 33% (360) were isolated directly from living or decaying plants (Harrison et al. 2024; Opulente et al. 2024). It is unknown why phytopathogens are relatively rare in the Saccharomycotina when so many are found in association with plants.

The most well-characterized plant-pathogenic Saccharomycotina species belong to the *Eremothecium* genus of the Saccharomycetales order. These fungi include *Eremothecium gossypii, Eremothecium ashbyi, Eremothecium coryli, Eremothecium sinecaudum*, *Eremothecium cymbalariae,* and *Eremothecium peggii,* which cause rot in a variety of plants, including citrus (Batra 1973; Crous et al. 2021), cotton bolls, coffee, soybean, tomato (Kurtzman et al. 2011), flax (Arnaud 1913), and mustard (Holley et al. 1984). The *Eremothecium* can cause significant crop damage (Batra 1973), yet relatively little is known about the mechanisms of pathogenesis in this group (Perez-Nadales et al. 2014). Oxylipin-covered ascospores in *E. sinecaudum* use a water-driven drilling movement to release spores—a mechanism that could facilitate plant infection (Leeuw et al. 2006). Additionally, decreased pathogenicity was observed in *Eremothecium* when oxylipin production was interrupted through exposure to aspirin (Leeuw et al. 2007).

Several other species in the Dipodascales order are known to be pathogenic to plants. *Geotrichum candidum* and *Geotrichum citri-aurantii*, commonly cause sour rot in crops, including citrus and tomatoes (Butler et al. 1988; Butler et al. 1965; Morris 1985; Wells 1977; Wild 1987). Similarly, *Geotrichum galactomycetum* has been reported to cause damage to tomatoes and lemons (Butler and Petersen 1972) and *Geotrichum reessii* has been reported as the agent of sour rot in tomatoes (Suwannarach et al. 2016). Other Saccharomycotina exhibit some qualities of plant pathogenesis but do not cause the level of losses to require widespread treatment with fungicides. *Kluyveromyces marxianus* (order Saccharomycetales) was observed to cause onion soft rot (Schroeder et al. 2007). *Botryozyma nematodophila,*(order Trigonopsidales) in association with the free-living nematode *Panagrellus zymosiphilus*, has been isolated from the sour-rot of grapes, but it is unclear if it is the causative agent of the infection as sour-rot is a complex ecological system (Hall et al. 2018; Kurtzman et al. 2011). Recent efforts have provided the genome sequences, functional annotations, and growth characterizations for nearly all known plant pathogenic species in the subphylum (Opulente et al. 2024) except for *E. ashbyi* and *E. peggii*. This new dataset allows us to identify genomic characteristics unique to Saccharomycotina plant pathogens.

Here, we utilize both a forward and a reverse ecology framework to identify genetic features characteristic of the plant-pathogenic Saccharomycotina. We first identified the genes believed to be involved in aspirin sensitivity in *Eremothecium.* These genes are related to 3-hydroxy oxylipin synthesis and were found to be broadly distributed across all the fungi with no notable differences in plant pathogenic fungi. Next, we conducted a pathway enrichment analysis to identify differences between plant-associated and plant-pathogenic fungi. This revealed that plant-pathogenic Saccharomycotina generally lacked enzymes required for rhamnose and nitrate metabolism. This result was surprising given the known role of these enzymes in defending fungi against cyanogenic glycosides.

## Results

### Diversity of plant-pathogenic and plant-associated Saccharomycotina

We divided the Saccharomycotina into 12 plant-pathogenic strains and 360 strains isolated directly from plants based on the yeast isolation environment ontology (Table S1; (Harrison et al. 2024)). We also leveraged the higher-level ecological categorizations from the ontology, which included plant, arthropod, chordate, environmental, and victual (food and drink) associations. All categorizations were made at the strain level. The taxonomic orders of plant-associated strains are shown in Figure 1. All twelve orders within the Saccharomycotina (Groenewald et al. 2023) had at least one plant-associated member, while the plant-pathogenic strains were found in three orders: Trigonopsidales, Dipodascales, and Saccharomycetales. At the species level (accounting for one species with two strains) there were four plant-pathogenic Dipodascales (*G. candidum, G. geotrichum, G. citri-aurantii, G. reessii*), one Trigonopsidales (*B. nematodophila*), and five Saccharomycetales (*E. gossypii*, *E. sinecaudum, E. coryli, E. cymbalariae,* and *K. marxianus*) species. These results highlight the broad taxonomic diversity among the plant-associated Saccharomycotina.

**Figure 1:**
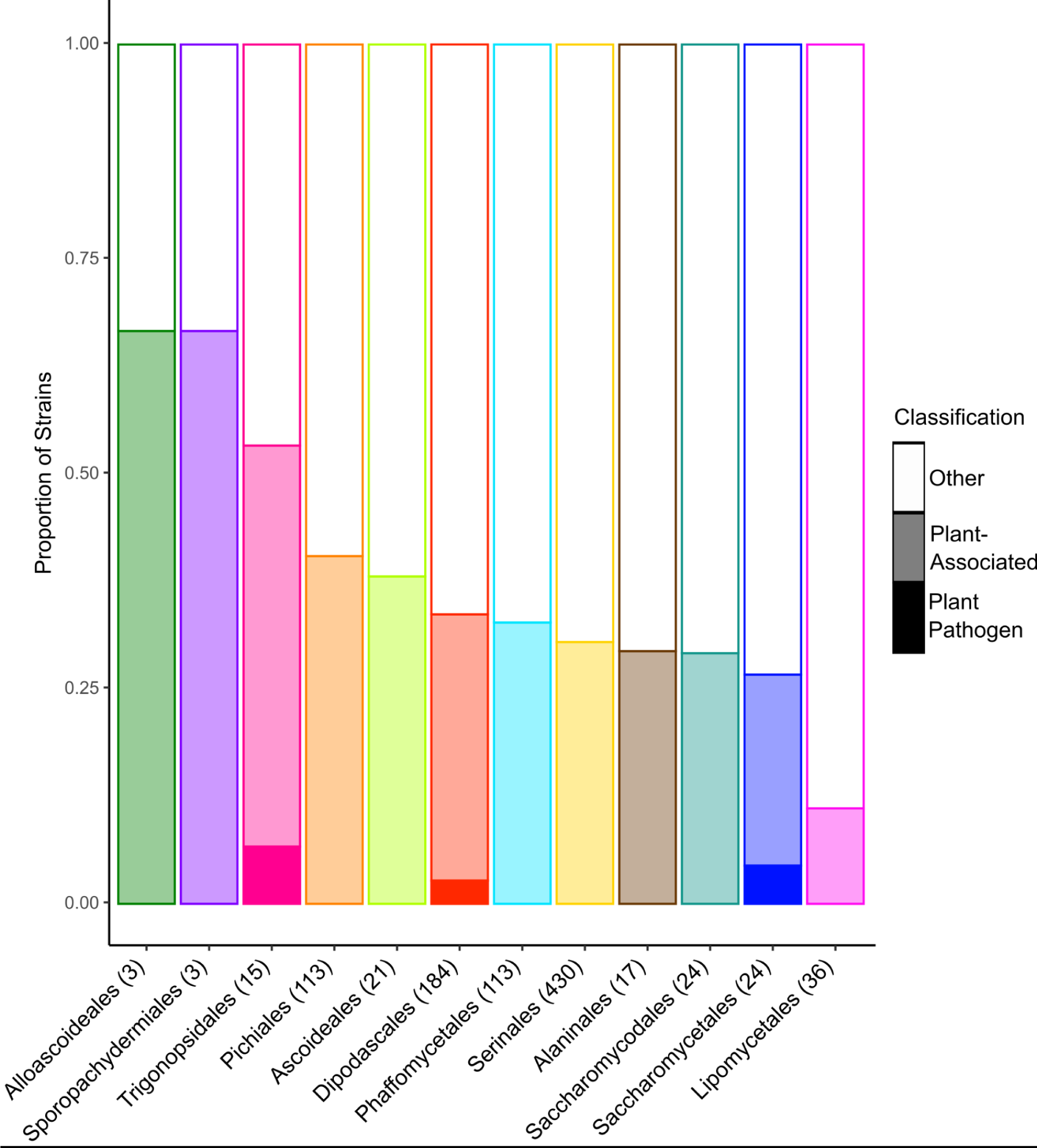
Taxonomic distribution of plant-associated and plant-pathogenic Saccharomycotina. All examined orders have strains that were isolated directly from plants. Conversely, the plant-pathogenic fungi were only found in three orders. The total number of strains within each order with known isolation environment information is shown in parentheses.

### 3-Hydroxy oxylipin synthesis

Oxylipins are important secondary metabolites frequently produced by fungi (Sebolai et al. 2012). In *Eremothecium*, 3-hydroxy oxylipins have a well-characterized role in ascospore release (Kock et al. 2003; Smith et al. 2000) and the sexual cycle (Leeuw et al. 2007). They are also believed to be involved in pathogenesis, potentially by facilitating invasion into plant tissue (Leeuw et al. 2006). The 3-hydroxy oxylipins are believed to be generated through incomplete β-oxidation of fatty acids in the Saccharomycotina (Sebolai et al. 2012). The enzymes that conduct β-oxidation are an acetyl-CoA dehydrogenase and an enoyl-CoA-hydratase (Sebolai et al. 2012). In *Saccharomyces cerevisiae,* the acetyl-CoA dehydrogenase is Pox1 (alias Fox1), and the enoyl-CoA-hydratase is Fox2. Most fungi conduct β-oxidation in the peroxisomes, as opposed to in the mitochondria (Poirier et al. 2006). Emerging evidence, however, suggests that some Saccharomycotina, such as *Clavispora lusitaniae* (syn. *Candida lusitaniae*), have a functional mitochondrial β-oxidation pathway (Gabriel et al. 2014). The mitochondrial and peroxisomal β-oxidation pathways both utilize the enzyme Fox2, suggesting enzymatic overlap between the peroxisomal and mitochondrial β-oxidation pathways. We investigated the presence of the acetyl-CoA dehydrogenase and enoyl-CoA-hydratase enzymes across the plant-associated and plant-pathogenic strains.

The acetyl-CoA dehydrogenase converts fatty acyl-CoA to trans-2-enoyl-CoA (Hiltunen et al. 2003). The *S. cerevisiae* acetyl-CoA dehydrogenase Pox1 belongs to the KEGG (Kyoto Encyclopedia of Genes and Genomes) Orthogroup (KO) K00232. We identified 2,213 genes in the KO K00232 across all the strains (Figshare Data). Of the 2,213 genes, we identified 23 putative horizontal gene transfer events of a gene from outside the Saccharomycotina subphylum based on the Pox1 gene tree structure and blast results (Figshare Data). The gene was not detected in 30 strains, including all 24 Saccharomycodales species in this study. The gene tree structure suggests a shared ancestry for the remaining 2,192 sequences with multiple duplication events. There were likely two duplication events in order Serinales, where 54% of species had three paralogs and 38% had two paralogs. Similarly, 37% of Dipodascales species had two paralogs of this gene, and 92% of Phaffomycetales had two or more paralogs. All plant-pathogenic Saccharomycotina, including the *Eremothecium,* had at least one acetyl-CoA dehydrogenase gene.

The enoyl-CoA-hydratase converts trans-2-enoyl-CoA to 3-ketoacyl-CoA and is known as Fox2 in *S. cerevisiae,* which maps to the KO K14729 (Hiltunen et al. 2003). We identified 1,159 genes in the KO (Figshare Data). Like the acetyl-CoA dehydrogenase, the Fox2 gene encoding the enoyl-CoA-hydratase was absent in 36 strains, including all the Saccharomycodales species sampled here. The gene sequences generally fell within their orders except for the Alaninales, which were nested within the Pichiales instead of as a sister clade. All plant-pathogenic strains, including the *Eremothecium,* had at least one enoyl-CoA-hydratase gene.

Based on gene presence and absence data, the oxylipin synthesis genes are not clearly associated with pathogenesis despite their role in *Eremothecium* development and aspirin sensitivity. Additional experiments and analysis will be needed to complete our understanding of how oxylipin synthesis enzymes contribute to Saccharomycotina plant pathogenesis

### Identification of pathways that differ between plant-associated and plant-pathogenic strains

To identify pathways that distinguish plant-associated from plant-pathogenic strains, we categorized KOs as present in 80% or more of the strains from each group and present in less than 20% of the strains from each group. We then performed a KEGG pathway enrichment analysis on the KEGGs present (>80%) and absent (<20%) in the plant-associated and plant-pathogenic strains (Table S2). These data were then filtered to identify pathways enriched in only one group. There were 13 pathways uniquely enriched in these categories (Table 2.) Most of these results (12 pathways) were unique to the plant-pathogenic strains. The two most significant pathways associated only with plant pathogens were “Fructose and mannose metabolism” and “Nitrogen metabolism.”

**Table 1:**
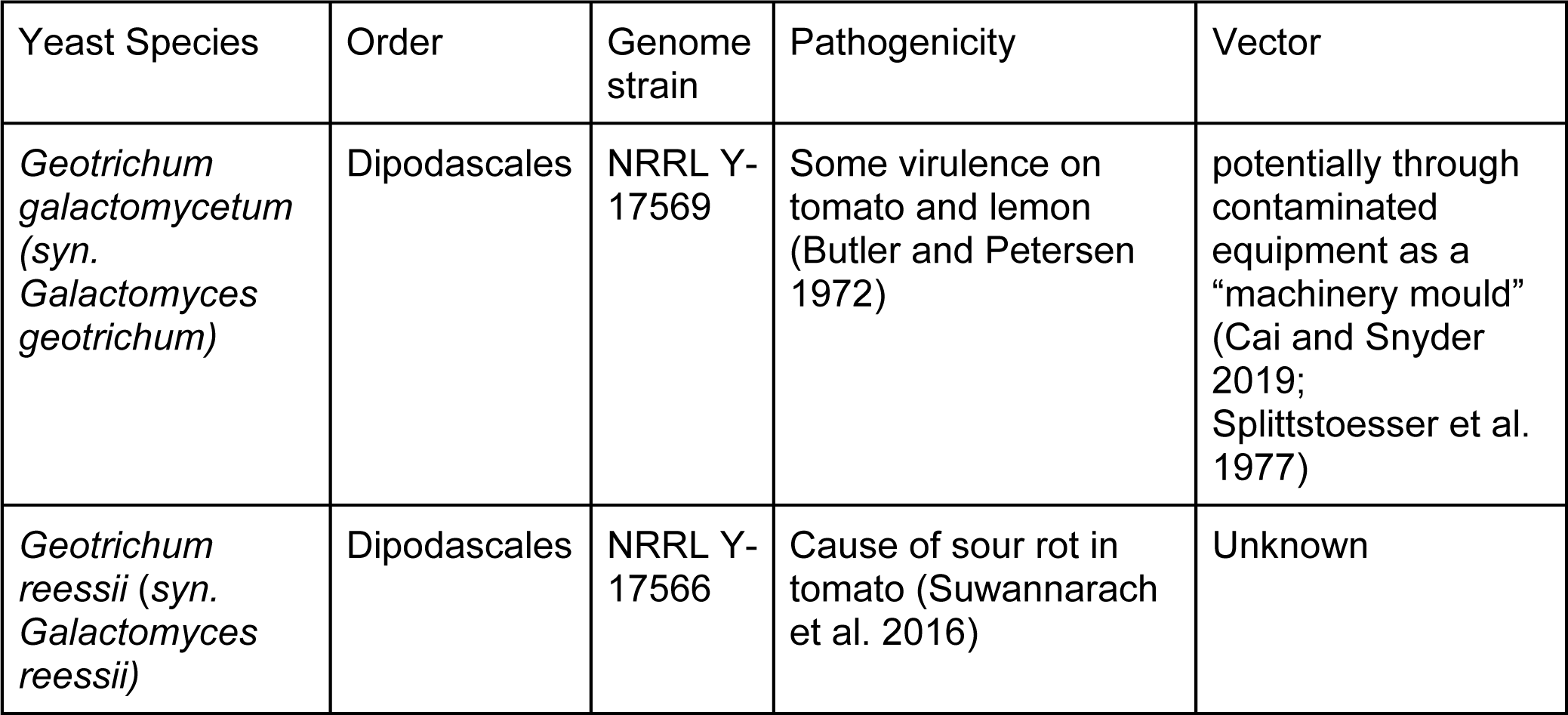
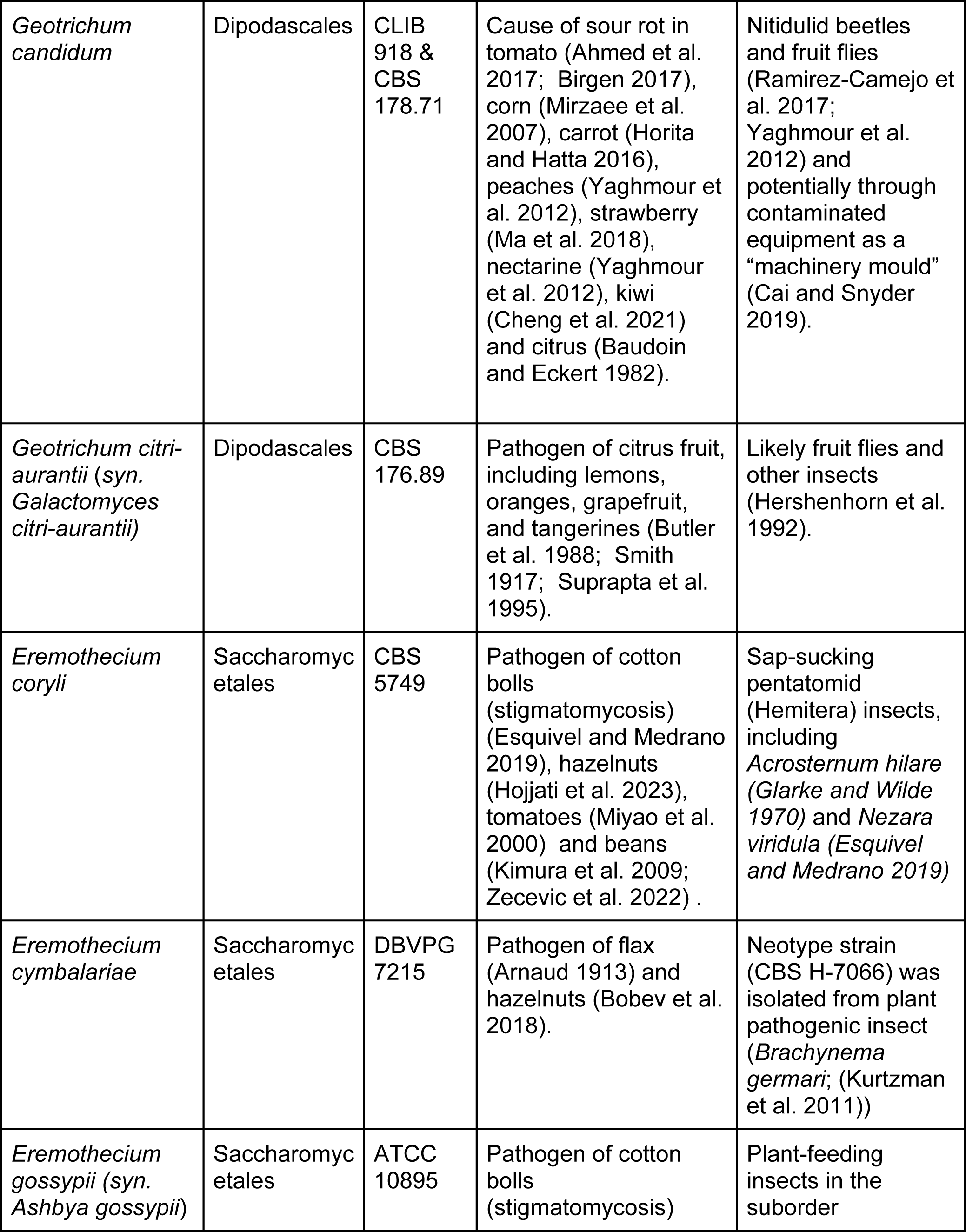
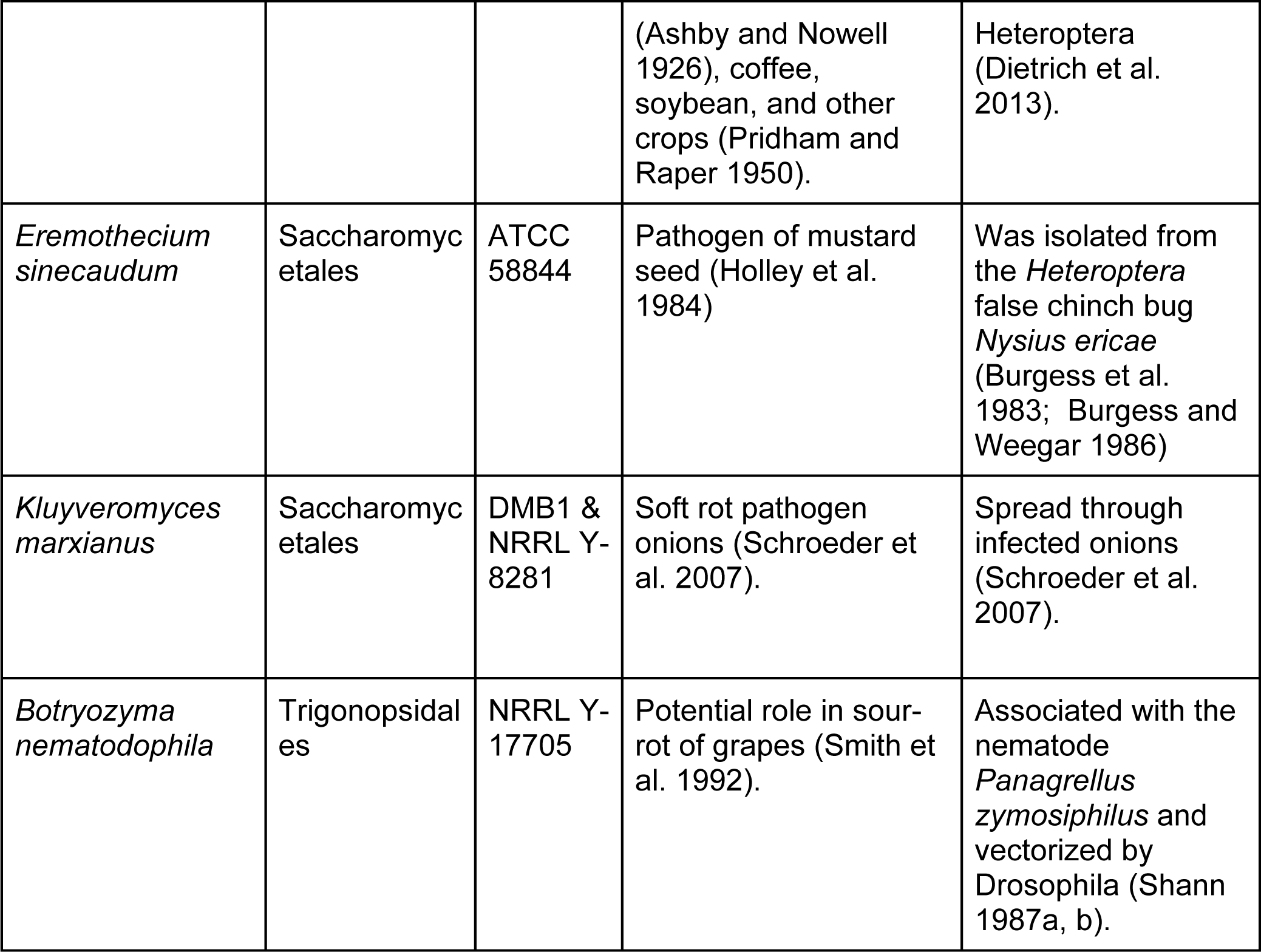
Saccharomycotina classified as plant pathogens. Based on the literature, we identified 10 species (12 strains total) of yeasts that are likely pathogens. The known plant host and possible vectors are also listed. These fungi primarily infect fruits and are vectorized by insects.

**Table 2:** KEGG pathway enrichment analysis of genes present or absent in plant-associated and plant-pathogenic Saccharomycotina. The most significant pathways identified in plant-pathogenic fungi that were not identified in plant-associated yeasts were fructose and mannose metabolism and nitrogen metabolism.

| Gene Category | Ecological Category | KEGG Pathway | Description | Adjusted P-value |
| --- | --- | --- | --- | --- |
| Absent | Plant-Associated | ko00280 | Valine, leucine and isoleucine degradation | 0.002 |
| **Absent** | **Pathogenic** | **ko00051** | **Fructose and mannose metabolism** | **0.002** |
| **Absent** | **Pathogenic** | **ko00910** | **Nitrogen metabolism** | **0.003** |
| Present | Plant-Associated | ko00563 | Glycosylphosphatidylinositol (GPI)-anchor biosynthesis | 0.009 |
| Absent | Pathogenic | ko00350 | Tyrosine metabolism | 0.020 |
| Present | Pathogenic | ko04110 | Cell cycle | 0.020 |
| Absent | Pathogenic | ko00052 | Galactose metabolism | 0.022 |
| Present | Pathogenic | ko00290 | Valine, leucine and isoleucine biosynthesis | 0.024 |
| Present | Pathogenic | ko04068 | FoxO signaling pathway | 0.033 |
| Absent | Pathogenic | ko02024 | Quorum sensing | 0.034 |
| Absent | Pathogenic | ko02020 | Two-component system | 0.035 |
| Absent | Pathogenic | ko01110 | Biosynthesis of secondary metabolites | 0.042 |

The fructose and mannose metabolism result was driven by the complete lack of rhamnose metabolism genes in the plant-pathogenic strains. Four enzymes convert L-rhamnose into L-lactaldehyde, and all of them are absent in the plant-pathogenic strains (Table S3). The four enzymes are L-rhamnonate dehydratase (K12661), L-rhamnose 1-dehydrogenase (K18337), 2-keto-3-deoxy-L-rhamnonate aldolase (K18339), and L-rhamnono-1,4-lactonase (K18338). These enzymes are found in 34%, 34%, 47%, and 33% of plant-associated strains, respectively. Overall, 31% (110/360) of the plant-associated strains had a complete L-rhamnose metabolism pathway compared to none of the plant-pathogenic strains.

Unsurprisingly, none of the plant pathogenic strains can grow on L-rhamnose when tested in the lab (Kurtzman et al. 2011; Opulente et al. 2024), while 16% (43/265) of the plant-associated strains with growth data were able to grow on L-rhamnose (Table S4). Across all strains measured, 15% of strains could grow on L-rhamnose. The lack of L-rhamnose metabolism genes is somewhat surprising, given that L-rhamnose is widely found in plants (Jiang et al. 2021). In plants, L-rhamnose is found in the cell wall and is used to make specialized metabolites, including glycoalkaloids (Jiang et al. 2021). In tomatoes and potatoes, L-rhamnose is needed to produce steroidal glycoalkaloids, which are used to defend against microbial and insect pests (McCue et al. 2007).

### Underrepresentation of nitrogen metabolism genes in plant-pathogenic strains

The nitrogen metabolism pathway was underrepresented in the plant-pathogenic but not the plant-associated fungi. Six KOs in the nitrogen metabolism pathway were completely absent or present in only one plant pathogenic strain but present in 14% or more of plant-associated strains (Table S5, Figure 2.) Each of these enzymes is involved in the processing of nitrogen-containing compounds into ammonia.

**Figure 2:**
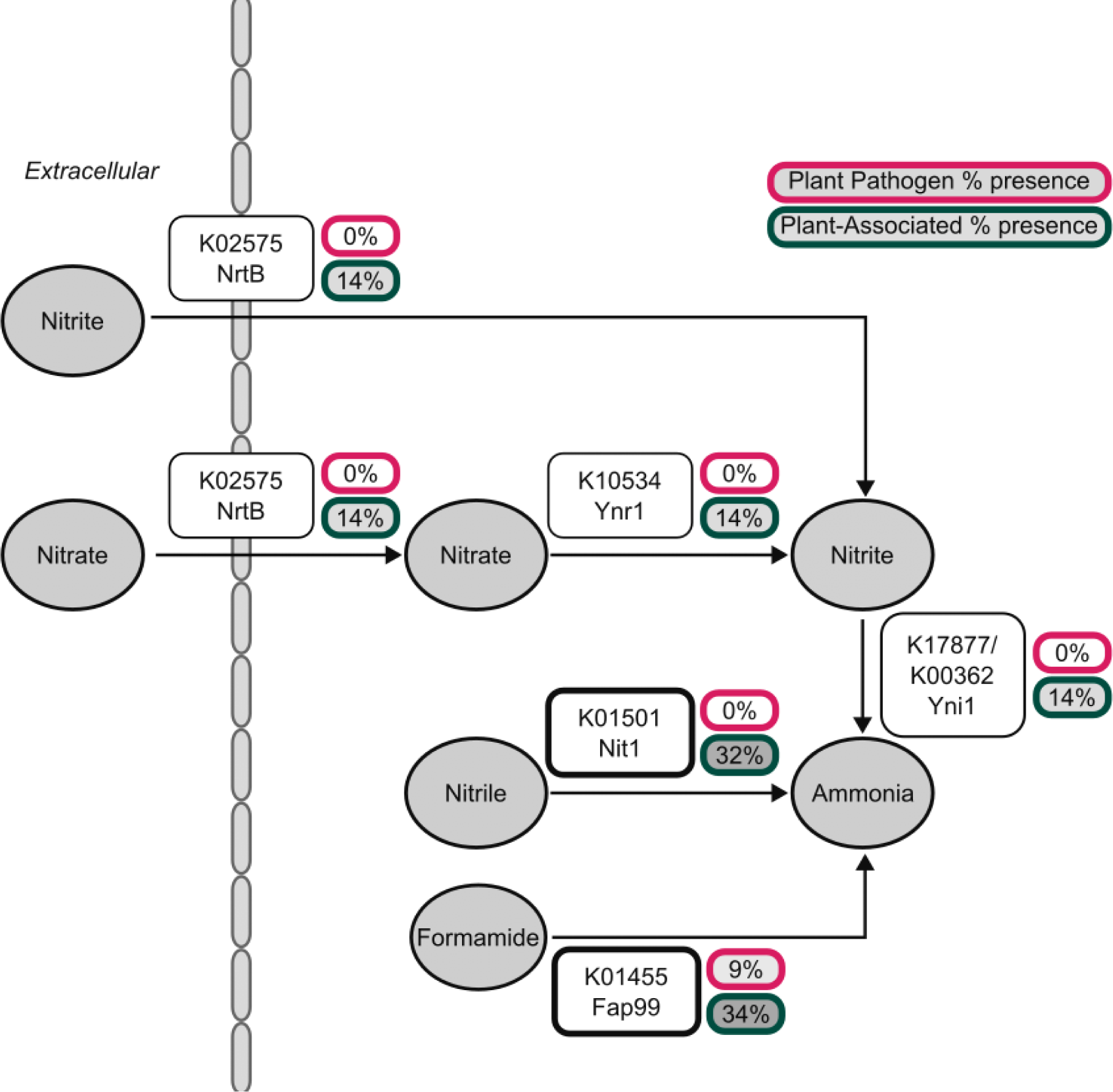
Presence of nitrogen metabolism genes identified in the pathway enrichment analysis. The plant-pathogenic strains almost entirely lack the necessary enzymes to metabolize nitrate, nitrile, or formamide into ammonia. This suggests that these strains cannot use nitrate or nitrite as their sole nitrogen source.

The nitrite reductase gene *YNI1*, previously characterized in *Ogataea polymorpha* (Siverio 2002), is associated with three KOs (K17877, K00362, and K00363). This gene is found in a gene cluster with the nitrate reductase gene *YNR1* (K10534) (Siverio 2002), which was also identified in this analysis. The other genes in this cluster are the nitrate transporter *YNT1* (K02575) and the transcription factor encoding genes *YNA1* and *YNA2* (no KEGG association.) We identified strains with putative nitrate assimilation gene clusters by identifying genomes with *YNI1*, *YNR1*, and *YNT1* KOs within a 10-gene window (Table S6, FigshareData.) We found that Saccharomycotina strains isolated from victuals (food or drink) had the highest proportion of genomes with the nitrate assimilation cluster (15 out of 92 genomes) and that Alaninales had the highest proportion (12 out of 17 genomes). In the 12 plant pathogen genomes, we identified no nitrate assimilation gene clusters. We also did not find a strong association between this cluster and the other environmental categories–the proportion of strains with this cluster ranges from 8% to 16% in arthropod, chordate, environmental, plant, and victuals categories. Given that this cluster is generally rare, we do not suspect this cluster’s absence contributes to plant pathogenicity.

We also identified a lack of the nitrate/nitrite transporter (K02575), encoded by *NRTB* in *O. polymorpha*, in the plant-pathogenic Saccharomycotina. Across all the strains, the transporter was found in 159 genomes, including all genomes predicted to have the nitrate/nitrite assimilation cluster (125 genomes.) The proportion of c gene is similar across environments, ranging from 11% in chordate-associated (8 of 74) to 17% in victuals-associated (16 of 92: Table S7.)

The KO K01455 was found in 1 plant pathogenic strain (*B. nematodophila*) and 34% (122/238) plant-associated strains. This KO is annotated as a formamidase and is associated with the gene *FAP99* in the human-pathogenic species *Candida albicans.* Generally, this enzyme hydrolyzes formamide into formic acid but has not been extensively studied in Saccharomycotina. It is known, however, that formamide is produced during cyanide degradation and that formamidases further metabolize formamide as a carbon or nitrogen source (Malmir et al. 2022). In the filamentous phytopathogenic fungus *Verticillium dahliae* the regulation of the formamidase gene is critical to pathogenicity (Xiao et al. 2024). The lack of formamidase in the majority of plant-pathogenic strains suggests they cannot use formamide as a nitrogen or carbon source and are unlikely to be able to metabolize cyanide produced by plants as a defense.

Finally, we analyzed the nitrilase (K01501), which was observed to be absent in the plant-pathogenic strains. This gene is characterized as *NIT1* in *S. cerevisiae.* The nitrilase KEGG annotation overlapped much less with the nitrate/nitrite assimilation cluster. Of the 369 genomes with a K01501 annotation, only 59 had the cluster. Across the orders, the K01501 annotation was found in all orders except for Alloascoideales, which has genomes available from three species (Table S8). The median presence was 33%, ranging from 13% in the Trigonopsidales (2 of 15) to 96% in the Saccharomycodales (23 of 24). The K01501 annotation was also widely distributed across strains isolated from different environments. It was found in approximately one-third of strains from all environments. The proportion of plant-associated strains with a nitrilase gene was 31% (114/360). This led to the hypothesis that the loss of nitrilases is a distinguishing feature of plant pathogens compared to plant-associated strains.

Interestingly, plant pathogenic and plant growth-promoting microorganisms have been shown to produce nitrilases (Barclay et al. 1998). Loss of nitrilases and other nitrilase superfamily enzymes, like cyanide hydratases and cyanide dihydratases, could result in a loss of cyanide and nitrile detoxification. For example, the rhizobacterium *Pseudomonas fluorescens* SBW25 produces a nitrilase, which allows it to tolerate a toxic level of nitriles produced by plants (Howden and Preston 2009). Similarly, in the filamentous plant pathogen *Fusarium solani* a cyanide hydratase allows this fungus to tolerate toxic levels of cyanide (Barclay et al. 1998). None of the Saccharomycotina, however, have enzymes mapped to cyanide hydratase (K10675) or cyanide dihydratase (K18282).

### Diversity of nitrilase genes in Saccharomycotina

In addition to *NIT1*, at least two other nitrilases are present in Saccharomycotina and are known as *NIT2* and *NIT3*. These enzymes belong to the nitrilase superfamily (Pace and Brenner 2001).To capture the full diversity of nitrilases in the subphylum, we conducted a thorough search of the genome annotations. We identified 3,963 putative nitrilase genes in an HMMer search using the KEGG annotated genes as the reference (Figshare Data). We then assigned these hits to previously calculated orthogroups (Opulente et al. 2024). Five distinct nitrilase orthogroups were identified: OG0003182 (748), OG0003106 (1143), OG000784 (1335), OG0004284 (550), and OG0004905 (186.) The orthogroups were characterized as follows by comparing them to the KEGG Data: OG0004284 to *NIT1* and K01501, OG0003106 to *NIT2* and K11206, OG000784 to *NIT3* and K13566. The orthogroup OG0004905 was also associated with K13566. OG0003182 corresponded to *NTA1* in *C. albicans* and K14663. The orthogroups OG0004284 (*NIT1*), OG0003106 (*NIT2*), and OG0000784 (*NIT3*) had much higher similarity to the nitrilase reference sequences (median e-values of 3.95e-83, 6.8e-81, and 9.4e-92 respectively) relative to OG0004905 and OG0003182 (median e-values of 1.15e-53 and 8e-59). Given the relatively higher e-values and associated KOs of OG0004905 and OG0003182, these were unlikely to be nitrilases and are not discussed further.

The pathway enrichment analysis identified the *NIT1* genes belonging to OG0004284. The comprehensive HMMER search identified an additional 85 *NIT1* instances in 73 species and did not include 9 previously identified instances in 4 species. However, the proportions of this gene in plant-pathogenic (0%: 0/12) and plant-associated (33% 120/360) species were not significantly different from the KO based analysis reported above (Table S9).

The gene *NIT2* encodes an amidase of deaminated glutathione involved in the metabolism and maintenance of glutathione (Peracchi et al. 2017), which is an antioxidant involved in plant development and stress responses (Bela et al. 2015). This enzyme was present in 88% of plant-associated (318/360) and 36% of plant-pathogenic strains (5/12; Table S9). The *NIT2* orthogroup was found in *K. marxianus*, *G. citri-aurantii*, *G. reessii, G. galactomycetum* and one of two stains of *G. candidum* (CLIB 918 and not CBS 178.71.) Glutathione plays a critical role in plant signaling and plant defense against pathogens (Dubreuil-Maurizi and Poinssot 2012). Deficiencies in plant glutathione synthesis enhance the susceptibility of *Arabidopsis thaliana* to fungal pathogens such as *Botrytis cinerea (Ferrari et al. 2003)*. Similar to NIT1, many of the plant pathogens, including the four *Eremothecium* species, lost the ability to process deaminated glutathione.

The gene *NIT3* (OG0000784) encodes an omega-amidase and was found in 73% (9/12) of the plant pathogenic strains and 90% (323/360) of the plant-associated strains (Table S9.) NIT3 may play a role in biofilm formation as the protein was identified in *C. albicans* biofilm extracts (Martinez et al. 2016) and secretomes (Vaz et al. 2021). These prior findings could indicate that Nit3 functions extracellularly, but this localization has not been demonstrated beyond model fungi.

## Discussion

It is counterintuitive that the loss of formamidase and nitrilase (*NIT1* and *NIT2*) genes were associated with plant-pathogenic Saccharomycotina. Formamidase and nitrilase genes have been shown to be involved in nitrogen assimilation and defense against plant compounds.

Studies in other plant pathogenic fungi suggest that nitrogen limitation occurs early in infection (Bolton and Thomma 2008) and that nitrogen utilization is key for virulent infection (Horst et al. 2012). Nitrilase superfamily enzymes in plant-associated bacteria may play a nitrogen assimilation role (Howden and Preston 2009). Plant-pathogenic strains, interestingly, rarely have the ability to assimilate nitrates in growth experiments (Kurtzman et al. 2011; Opulente et al. 2024). We found no plant-pathogenic strains that could assimilate nitrate, while 14% (48/337) of the plant-associated strains had the ability. These results suggest that paradigms developed around nitrogen assimilation in other Ascomycota plant pathogenic fungi are unlikely to be applicable in Saccharomycotina.

Plants also use nitrile-containing secondary metabolites as defense mechanisms. For example, cyanogenic glycosides, a type of α-hydroxynitrile, defend against insect herbivores (Gleadow and Moller 2014). Plants also use phenylacetonitrile for defense (Yactayo-Chang et al. 2020). Insects ingest these nitriles and break them down into cyanide, which kills them (Martinez and Diaz 2024). Fungi are known to utilize nitrilase superfamily enzymes to degrade cyanide; this includes species such as *Neurospora crassa, Aspergillus nidulans, Fusarium graminearum* (Basile et al. 2008) and *Gloeocercospora sorghi* (Wang et al. 1992). The Saccharomycotina plant-pathogenic strains lack nitrilase enzymes to defend against these compounds.

Multiple hypotheses could explain why the plant-pathogenic Saccharomycotina have a surprising lack of nitrilase genes, which have been previously associated with other fungal plant pathogens. One hypothesis is that their vectors have shaped Saccharomycotina plant pathogen evolution. This is the case in the well-studied yeast-cactus-*Drosophila* system, where the three organisms exhibit coadaptation (Starmer and Fogleman 1986). The *Eremothecium* are known or theorized to be spread by insects that pierce the fruit and facilitate infection. (Table 1). Additionally, *G. candidum*, *G. citri-aurantii*, and *B. nematodophila* are likely spread by insects such as fruit flies. Fruit flies can puncture fruit surfaces to deposit larvae which suggests they can vectorize plant fungal pathogens. For example, the invasive *Drosophila suzukii* has been associated with the spread of fruit rot pathogens such as *B. cinerea* in strawberries (Lewis et al. 2019). In these cases, the Saccharomycotina rely on the vector to bypass the physical barriers outside of fruits. This reliance on vectors for invasion of plant tissue may have limited the evolution of the Saccharomycotina plant pathogens.

Similarly, the Saccharomycotina plant pathogens may have evolved to avoid cyanogenic glycosides. Cotton, a major host for *E. gossypii*, produces secondary metabolites as a defense mechanism, such as terpenes, but is not known to produce cyanogenic glycosides (Stipanovic et al. 2010). Similarly, *G. candidum* causes sour rot in citrus plants, which are also not known to produce cyanogenic glycosides (Ali et al. 2014). This pattern suggests that the presence of cyanogenic glycosides and cyanide, even in small amounts, may prevent the Saccharomycotina plant pathogens from colonizing plant tissue. This is also consistent with the observation that Saccharomycotina pathogens target fruits as opposed to other plant parts like leaves, stems, or roots. Fruit defenses are highly complex and vary during the transition from immature to mature and dispersed fruits (Whitehead et al. 2022). During ripening, fruits undergo morphological and biochemical changes, such as cell wall degradation and changes in phytohormone (including aspirin or salicylic acid) production, that can result in increased susceptibility to fungal infection (Alken and Fortes 2015). For example, immature strawberries contain sufficient proanthocyanidins to prevent *B. cinerea* infection, but as the strawberry ripens, the activity of the proanthocyanidins decreases, allowing the fungus to infect the fruit (Jersch et al. 1989). Overall, Saccharomycotina appear to be limited by their vectors and genetic composition to infecting ripening fruits.

## Materials & Methods

### Data source

Genomes, annotation, isolation environment, and phylogeny for 1,154 Saccharomycotina strains were obtained from published work (Opulente et al. 2024). Genera for which no living culture was available or those described after February 2021 were not included in Opulente et al. (2024) and, therefore, not included in our study. Species names (Supplemental Table 1) are accurate as of February 2021 except for the 12 plant-pathogenic strains (Table 1) which have been updated as of February 2025. We defined plant-associated strains as those isolated directly from plants as defined by the Ontology of Yeast Environments (OYE) (Supplemental Table 1, (Harrison et al. 2024).) Strains pathogenic to plants were defined from the literature. Strains characterized as plant pathogens and their associated references are shown in Table 1.

### Gene enrichment analysis

We conducted a gene enrichment analysis to identify the pathways and modules enriched or depleted in the plant-pathogenic as compared to plant-associated strains. Kyoto Encyclopedia of Genes and Genomes (KEGG) Orthologs (Kanehisa and Goto 2000) in each ecological group were split into those present in >80% of the genomes and those present in <20% of the genomes (Figshare Data.) The R package clusterProfiler v 4.101.1 (Yu et al. 2012) was used to find enriched and depleted genes in each category using all KEGG annotations present in any of the 1,154 genomes as the possible universe. We removed any pathways associated with “Human Diseases” or “Organismal Systems” as they have limited applicability to single-celled fungi. P-values were corrected for multiple testing using false discovery rate correction. Pathways that were enriched or depleted in both the plant-associated or plant-pathogen were filtered from the analysis.

### Gene identification methods

For several genes of interest, we further refined the annotation. We identified reference sequences for these genes using either NCBI protein BLAST results or the genes identified in the previous KEGG annotation (Opulente et al. 2024). The reference proteins (Figshare Data) were used to generate profile HMMs using the HMMer v3.3.2 (hmmer.ogr) package. We found amino acid sequences in the genome annotations with HMMERSearch with E-values less than 1E-50 and the profile HMM. Finally, we compared these results to the previously identified orthogroups (Opulente et al. 2024). Combining sequence similarity (KEGG and HMMer) and evolutionary information (orthogroups) allowed us to confidently identify the genes within each genome.

We also built gene trees to visually inspect the evolution of the specific genes of interest. Amino acid sequences were aligned using MAFFT v7.273 (Katoh et al. 2002) and gene trees were constructed using IQ-Tree v2.1.2 (Minh et al. 2020) which included the identification of gene models. Trees were visualized in iTOL (Letunic and Bork 2021).

## Acknowledgements

We thank Dr. Morgan Carter (UNC Charlotte) and members of the Y1000+ Project for their feedback on the work. Computational analyses were run in the UNC Charlotte high performance computing cluster in Charlotte, North Carolina.

## Supplemental Data

All supplemental data is currently hosted on the FigShare repository.

Table S1: Strain designation, genome information and taxonomic order for all 1,154 strains in the dataset. This table also includes the environment from which the strain was isolated and the categorization of that environment.

Table S2: KEGG pathway enrichment results. The unfiltered results for the KEGG pathway enrichment analysis, including all pathways enriched in both plant-associated and plant-pathogenic strains.

Table S3: Presence and absence of rhamnose metabolism genes in plant-associated and plant-pathogenic strains. The strain count and strain percents are shown.

Table S4: Binary growth of strains on rhamnose and associated citations

Table S5: Presence and absence of nitrilase genes in the plant-associated and plant-pathogenic strains. This includes the raw strain count and percentages.

Table S6: Presence of one or more nitrate reductase clusters in the Saccharomycotina.

Table S7: Strains with the nitrate/nitrite transporter (K02575), encoded by *NRTB* in *O. polymorpha*.

Table S8: Presence and absence of the nitrilase (K01501) gene across the strains.

Table S9: Presence of characterized orthogroups in the plant-associated and plant-pathogenic fungi.

